# Activity or Connectivity? Evaluating neurofeedback training in Huntington’s disease

**DOI:** 10.1101/481903

**Authors:** Marina Papoutsi, Joerg Magerkurth, Oliver Josephs, Sophia E Pépés, Temi Ibitoye, Ralf Reilmann, Nigel Hunt, Edwin Payne, Nikolaus Weiskopf, Douglas Langbehn, Geraint Rees, Sarah J Tabrizi

**Affiliations:** UCL Huntington’s Disease Centre, Queen Square Institute of Neurology, University College London, UK; Birkbeck-UCL Centre for Neuroimaging, University College London, London, UK; Wellcome Centre for Human Neuroimaging, Queen Square Institute of Neurology, University College London, UK; University of Oxford, UK; George Huntington Institute and Dept. of Radiology University of Muenster, Germany; Section for Neurodegeneration and Hertie Institute for Clinical Brain Research, University of Tuebingen, Germany; Eastman Dental Institute, University College London, UK; Max Planck Institute for Human Cognitive and Brain Sciences, Leipzig, Germany; Carver College of Medicine, University of Iowa, USA; Institute of Cognitive Neuroscience, University College London, UK; UK Dementia Research Institute at University College London, UK

**Keywords:** Neurofeedback training, neuroplasticity, Huntington’s disease, real-time fMRI

## Abstract

Non-invasive methods, such as neurofeedback training (NFT), could support cognitive symptom management in Huntington’s disease (HD) by targeting brain regions whose function is impaired. The aim of our single-blind, sham-controlled study was to collect rigorous evidence regarding the feasibility of NFT in HD by examining two different methods, activity and connectivity real-time fMRI NFT. Thirty-two HD gene-carriers completed 16 runs of NFT training, using an optimized real-time fMRI protocol. Participants were randomized into four groups, two treatment groups, one receiving neurofeedback derived from the activity of the Supplementary Motor Area (SMA), and another receiving neurofeedback based on the correlation of SMA and left striatum activity (connectivity NFT), and two sham control groups, matched to each of the treatment groups. We examined differences between the groups during NFT training sessions and after training at follow-up sessions. Transfer of training was measured by measuring the participants’ ability to upregulate NFT target levels without feedback (near transfer), as well as by examining change in objective, a-priori defined, behavioural measures of cognitive and psychomotor function (far transfer) before and at 2 months after training. We found that the treatment group had significantly higher NFT target levels during the training sessions compared to the control group. However, we did not find robust evidence of better transfer in the treatment group compared to controls, or a difference between the two NFT methods. We also did not find evidence in support of a relationship between change in cognitive and psychomotor function and NFT learning success. We conclude that although there is evidence that NFT can be used to guide participants to regulate the activity and connectivity of specific regions in the brain, evidence regarding transfer of learning and clinical benefit was not robust. Although the intervention is non-invasive, given the costs and absence of reliable evidence of clinical benefit, we cannot recommend real-time fMRI NFT as a potential intervention in HD.

## Introduction

Neurofeedback training (NFT) is a non-invasive intervention used to train participants in a closed-loop design to regulate their own brain activity(Sitaram *et al.*, 2017). The underlying principle is that by regulating different aspects of their brain activity, e.g. regional activation or inter-regional connectivity, participants would implicitly regulate associated cognitive function. Huntington’s disease (HD) is a genetic neurodegenerative condition characterised by progressive motor, psychiatric and cognitive impairment, as well as early striatal atrophy, cortical and cortico-striatal connectivity loss(Tabrizi *et al.*, 2011; Poudel *et al.*, 2014; McColgan *et al.*, 2015; Novak *et al.*, 2015). There are currently no treatments for cognitive impairment in HD and the effect of disease-modifying therapies, such as antisense-oligonucleotide approaches (ASO (Tabrizi *et al.*, 2019*b*)), on cognitive function is, at present, unknown. Our motivation for testing NFT, is that it, if successful, it could be used as an adjunct treatment to invasive, disease-modifying therapies(Linden and Turner, 2016, Tabrizi *et al.*, 2019*a*). However, there are several challenges in designing effective NFT trials and testing their efficacy, including the choice of an appropriate NFT target for the specified clinical population.

Because striatal atrophy and cortico-striatal connectivity loss appear early on in HD and correlate with cognitive and psychomotor impairment(Tabrizi *et al.*, 2009, 2011; Poudel *et al.*, 2014; McColgan *et al.*, 2015; Novak *et al.*, 2015), striatal activity and cortico-striatal connectivity would be the obvious targets for NFT. NFT could therefore be used to “boost” the activity or connectivity of the striatum in HD gene-carriers at pre-symptomatic or early stages of the disease, i.e. while levels of atrophy are still low. In a recent proof-of-concept study we used the supplementary motor area (SMA) as a target for real-time fMRI NFT in HD patients(Papoutsi *et al.*, 2018). We selected BOLD fMRI signal from the SMA because it can be reliably measured in real-time(Subramanian *et al.*, 2011, 2016), and its function and connectivity to the striatum is disrupted by HD(Klöppel *et al.*, 2009). Previous studies have also shown that NFT induced changes are not just localised to the target region, but extend to a wider network of regions(Horovitz *et al.*, 2010; Ruiz *et al.*, 2013; Emmert *et al.*, 2016), suggesting that a proxy region would be appropriate. We found that HD patients can be trained to increase the level of SMA activity and that improvement in cognitive and psychomotor behaviour after training related to increases in activity of the left Putamen and SMA – left Putamen connectivity during training. This suggested that SMA-striatum connectivity could be a more appropriate NFT target than SMA activity in HD.

The aim of the current study is to compare the two NFT approaches, SMA activity and SMA-striatum connectivity, and to collect rigorous evidence on the feasibility of the method in HD. We used BOLD fMRI signal change from the SMA as the target for activity NFT and correlation between the signal from the SMA and left striatum during upregulation as the target for connectivity NFT(Megumi *et al.*, 2015; Yamashita *et al.*, 2017). In addition, we used a single-blind, randomized, sham-controlled design and employed an optimized real-time fMRI processing pipeline using a prospective-motion correction system (PMCS; (Zaitsev *et al.*, 2006; Todd *et al.*, 2015) for real-time head motion correction and real-time physiological noise filtering(Misaki *et al.*, 2015) to ensure that we obtain high quality evidence. Finally, to ensure participant blinding and control for experimental exposure and motivation, participants that were randomized to the control group were yoked to a participant in the treatment group, and received feedback based on the NFT target levels of their yoked participant from the treatment group, rather than their own(Thibault *et al.*, 2016; Sorger *et al.*, 2019). This setup enables us to collect high quality evidence regarding the use of real-time fMRI NFT for the treatment of cognitive impairment in HD.

## Materials and Methods

### Participants

Thirty-four adults with HTT gene CAG expansion greater than 40 were recruited to the study. One participant withdrew from the study and another participant was excluded, because a large number of trials were contaminated with motion-related artifacts. Details on the remaining thirty-two participants who were included in the analyses are shown in Table 1. There were no statistically significant differences between the treatment and control groups for the two types of NFT (using a non-parametric Mann-Whitney test all p > 0.2). All participants provided written informed consent according to the Declaration of Helsinki and the study was approved by the Queen Square Research Ethics Committee (05Q051274).

**Table 1:**
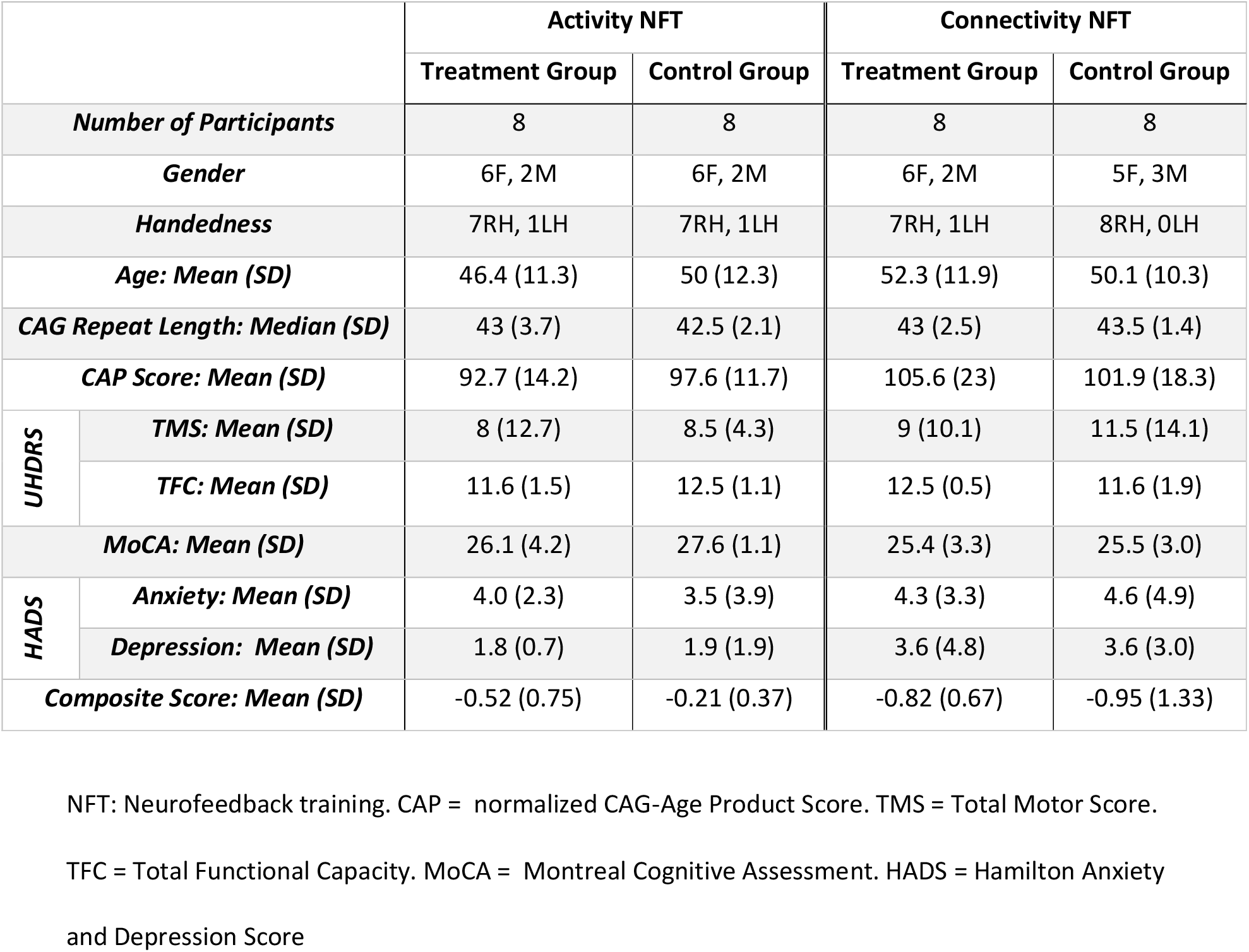
Demographic Information

Information regarding sample size calculations prior to the start of the study are provided in the supplementary materials. Briefly, the present study was powered in order to be able to detect, previously reported, very large differences (Cohen’s d effect size = 1.65 and 1.60; see supplementary materials) between treatment and sham NFT control groups in near transfer. As this was a feasibility study, we chose to power on near transfer and not far transfer effects. Although near transfer effects are not clinically relevant, they do allow us to test NFT learning transfer and as such can serve as a suitable endpoint for this feasibility study. If the findings from this study are promising, then the effect sizes estimated from this study could be used to power a future RCT focusing on efficacy.

### Study Structure

As part of the study, participants completed 1 screening, 1 baseline, 4 neurofeedback training and 3 follow-up sessions. The first follow-up was within 2 weeks from the last training visit, the second between 4-6 weeks and the third between 8 and 10 weeks (also see Supplementary Table 2). A diagram of the study design is shown in Figure 1. A Prospective Motion Correction system (PMCS) was used to correct head motion during scan acquisition(Zaitsev *et al.*, 2006; Todd *et al.*, 2015). Details of the PMCS are provided in the supplementary materials. Participants who consented to the use of the PMCS had teeth impressions acquired during the screening visit by a qualified orthodontist (NH). During the screening session participants completed the Montreal Cognitive Assessment test (MoCA(Nasreddine *et al.*, 2005) and a number of cognitive and psychomotor tasks. The purpose of the testing on the screening visit was to familiarize the participants with the tests and minimize practice effects during follow-up. The same measurements were repeated during the baseline and follow-up sessions.

**Figure 1:**
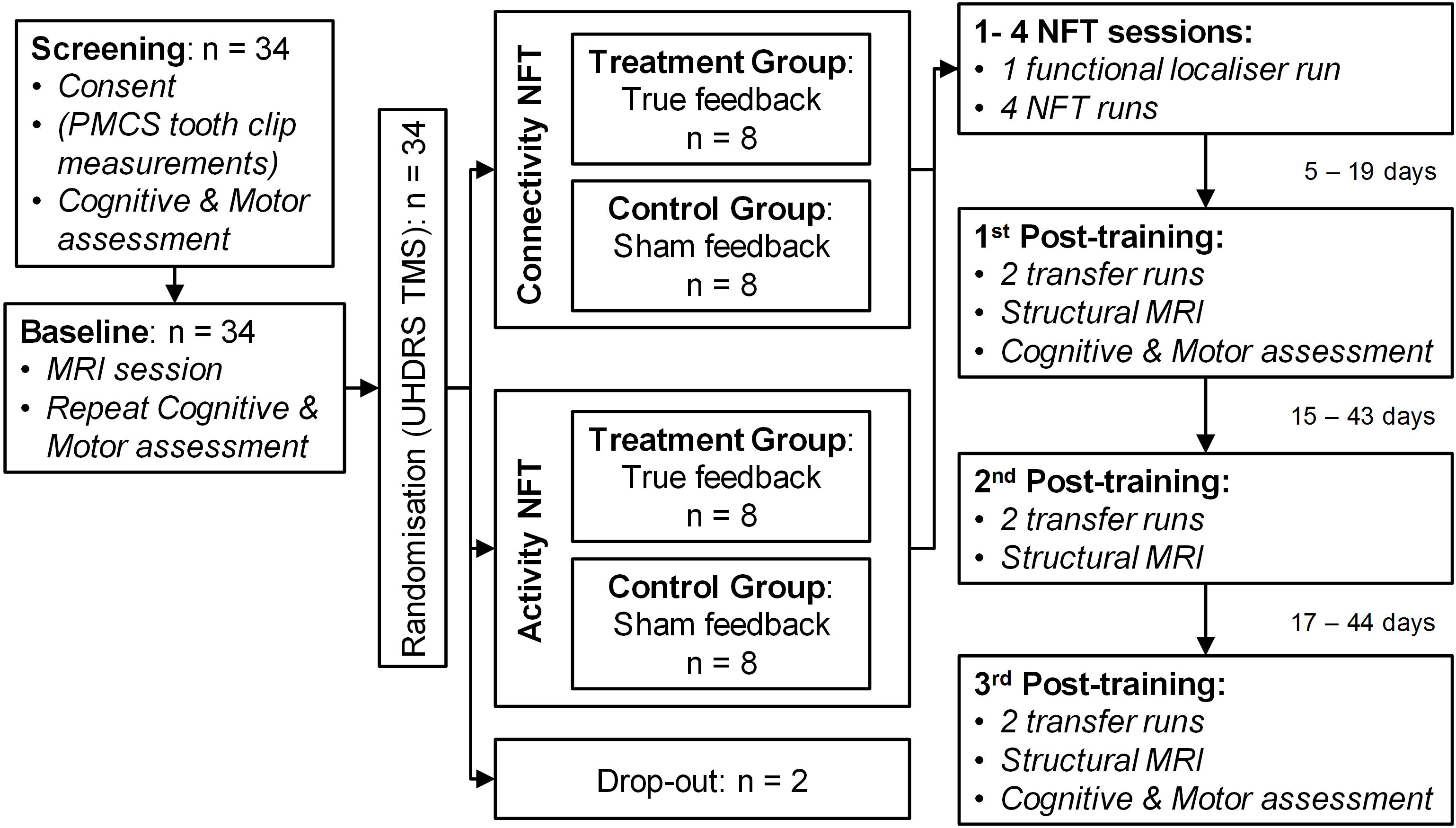
Diagram of study structure. PMCS = Prospective motion correction system. UHDRS TMS = Unified Huntington’s disease Rating Scale Total Motor Score. NFT = Neurofeedback training.

To assess change in cognitive and psychomotor function following NFT we calculated a composite score using the same procedure and measures as in our previous study(Papoutsi *et al.*, 2018). In summary, these measures were selected a-priori based on previous work showing that they are sensitive to disease progression(ref), they were converted to z-scores and summed to create the composite score. The measurements included were: number correct for Stroop Word Reading, number correct for Symbol Digit Modalities Test (SDMT), annulus length for Indirect Circle Tracing (log transformed), number correct for negative Emotion Recognition, inter-tap interval and standard deviation of inter-onset interval (log transformed) during speeded tapping with the non-dominant index finger, and standard deviation of mid-tap interval deviation from target rhythm (log transformed) for paced tapping with the non-dominant index finger at 1.8Hz.

The baseline and follow-up sessions included: 1) repetition of the cognitive and psychomotor testing (only on the first and third follow-up), 2) structural MRI measurements and 3) two fMRI runs assessing the participant’s ability to upregulate the target NFT measures without neurofeedback. The fMRI runs consisted of 5 upregulation blocks (30s each), 6 rest blocks (30s each) and 5 response blocks (18s each; see Supplementary Figure 2B). Similar to our previous study(Papoutsi *et al.*, 2018) we used a simple attention task during the rest blocks, whereby participants monitored changes in the luminance of a white bar. After the baseline session participants were randomized to one of four groups: activity NFT treatment and control groups, and connectivity NFT treatment and control groups. Randomization was based on the Unified Huntington’s Disease Rating Scale(Huntington Study Group, 1996) (UHDRS) Total Motor Score (TMS). More details regarding the randomization procedure are provided in the supplementary materials.

NFT sessions started with a fist-clenching run used to select the target ROIs. Participants were instructed to clench their left fist during the active blocks (10 blocks, 20.4s duration) and rest during the rest blocks (11 blocks lasting 20.4s each; see Supplementary Figure 2A). Using Turbo-BrainVoyager (TBV; Brain Innovation, The Netherlands) the fMRI run was analysed in real-time and the resulting statistical map was used to define the ROIs for the subsequent NFT runs. Participants completed 4 NFT sessions on different days and each session included 4 NFT runs (two participants completed 3 runs on one of the NFT sessions, because of fatigue). The activity NFT runs consisted of 6 rest blocks (30s duration), 5 response blocks (18s) and 5 upregulation blocks (30s; see Supplementary Figure 2C). The rest and response blocks were identical to those of the transfer runs described previously. During the upregulation blocks feedback was presented continuously in the form of a red bar. In the treatment group the height of the red bar represented the percent signal change at a given point during the upregulation block vs the mean activation during the preceding rest block. Once the upregulation blocks started, there was an average delay of 2s until the red bar appeared and then it was updated every 1.2s. The connectivity NFT runs consisted of 5 rest blocks (45s), 5 upregulations blocks (30s) and 5 feedback blocks (3s; see Supplementary Figure 2D). Feedback was presented intermittently at the end of the upregulation blocks in the form of a red bar. In the treatment group the height of the red bar was calculated using the Pearson’s correlation coefficient between the SMA and left striatum ROI time-series during the upregulation blocks only(Megumi *et al.*, 2015). In both cases (activity and connectivity NFT) the feedback provided to the sham control groups was calculated using data from a yoked participant in the corresponding treatment group. More details on the sham neurofeedback setup and the real-time fMRI setup are provided in the following paragraphs.

Similar to our previous study, we used shaping in both cases in order to facilitate learning and motivation(Weiskopf *et al.*, 2004; Linden *et al.*, 2012; Papoutsi *et al.*, 2018), whereby the difficulty in increasing the height of the feedback bar was adjusted according to the participants’ performance in the preceding block.

### Target ROI selection

The NFT target ROIs were drawn at the start of each NFT session using TBV. For the activity NFT sessions, the SMA was selected as the target ROI. For the connectivity NFT sessions, the SMA and the left striatum (including putamen, globus pallidus and caudate) were selected as the target ROIs. Similar to our previous study(Papoutsi *et al.*, 2018) and comparable to other studies(Subramanian *et al.*, 2011; Paret *et al.*, 2014, 2016; Nicholson *et al.*, 2017), the ROIs were re-drawn at each session ensuring that only voxels with high activation are selected. The ROIs from the first visit were used as a reference, when drawing the ROIs for the subsequent visits to ensure that the position was similar, although the exact voxels selected might be different. For the SMA the statistical map was thresholded at t-value = 3 and a rectangle was drawn around the SMA cluster for the active vs rest contrast. The location of the striatum was identified visually on the first EPI scan of the localiser run using landmarks and the EPI contrast. Due to high iron concentration, the putamen and globus pallidus appear darker on an EPI scan and are therefore easy to identify on EPI scans. A rectangle was drawn around the striatum including the putamen, globus pallidus, caudate and ventral striatum. Because of the rectangular shape, the striatal ROI, also included surrounding white matter. However, the ROI was centred around the striatum and most of the recorded signal originated from the gray matter of the striatum. A heat map showing the overlap of the ROIs across all participants is shown in Figure 2A and B. We chose to define our ROIs using a functional localiser and anatomical landmarks, rather than creating an anatomical mask, because it was not always possible to acquire a structural MRI volume during the baseline visit due to patient fatigue.

**Figure 2:**
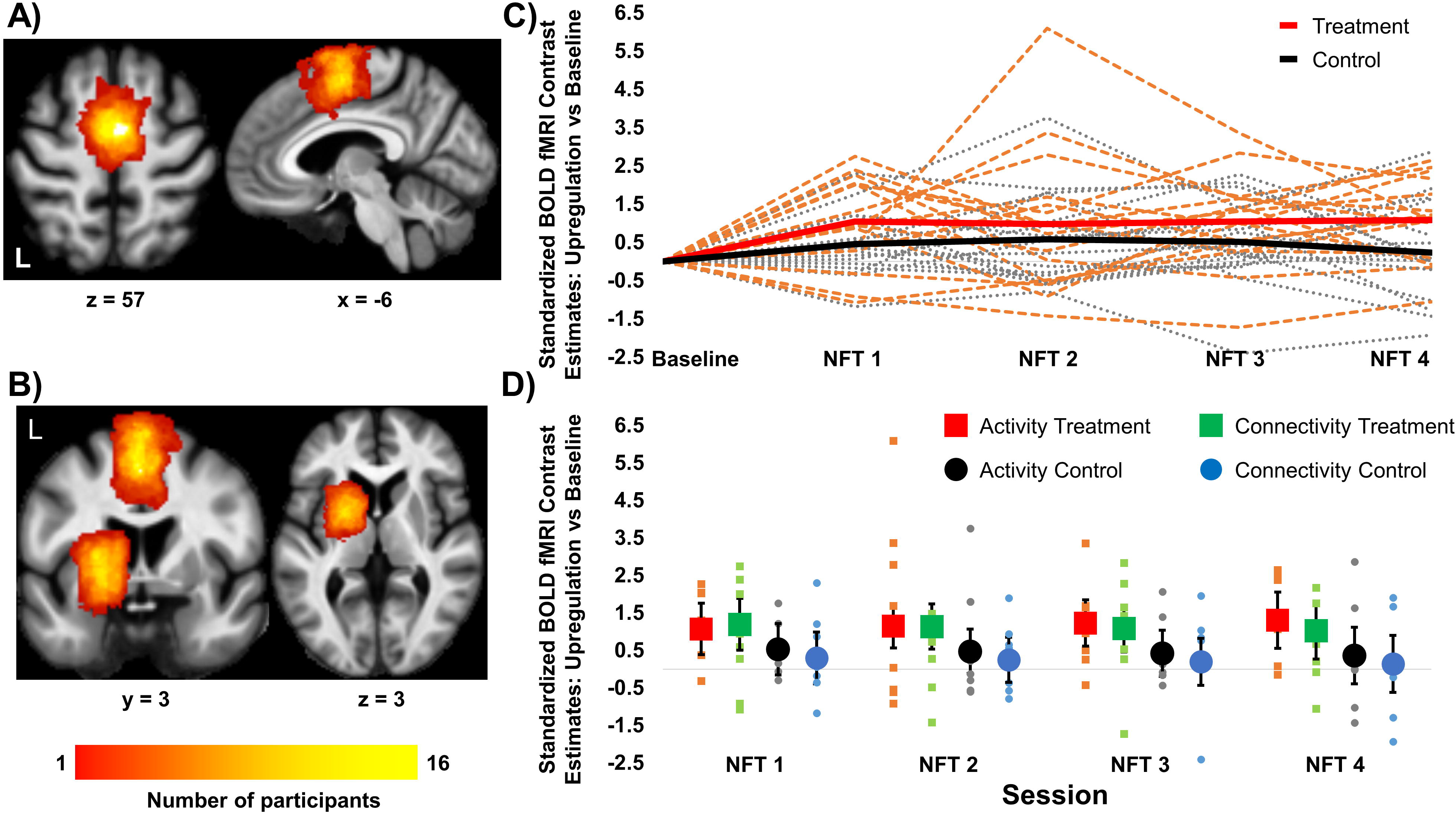
Learning effects in activity and connectivity NFT. (A) and (B) Heat maps showing the location and overlap of the target ROI across all participants in the activity and connectivity NFT groups respectively. Maps are superimposed on a group average MT image. (C) Change from baseline in the target NFT levels across all training sessions per subject (dotted lines). The group mean per session is shown with thick continuous lines. Shown in red (group mean) and orange (individual participants) is the treatment group, whereas shown in black (group mean) and gray (individuals) is the control group collapsed across both types of NFT. (D) Dot plots show the change in NFT target levels from baseline across all NFT sessions for the four subgroups: activity treatment group (red circles), connectivity treatment group (green circles), activity control group (black squares) and connectivity control group (blue squares). The horizontal gray lines in the dot plots show the baseline, data points above this line represent an increase compared to baseline. The small squares and circles are the individual data points, whereas the larger squares and circles show the adjusted mean group effects. Error bars are 95% CI.

### Sham Neurofeedback

We chose to use a sham neurofeedback for the control groups in order to control for potential placebo effects as a result of recruiting participants to an interventional study(Foroughi *et al.*, 2016; Thibault *et al.*, 2016; Sorger *et al.*, 2019). By choosing the “yoked” approach we ensured that the feedback control participants received was biologically plausible and matched to that of the treatment group. We chose not to use the approach of using a different ROI for the control group, because of potential problems with the spread of training effects across other brain regions. We do not yet understand the mechanism underlying NFT in HD and how widespread any effects could be, therefore we were not certain which other regions in the brain would be appropriate to use as control targets(Mehler *et al.*, 2018). To confirm whether the feedback received by the participants was contingent to their own brain activity, we performed confirmatory analyses after the end of the study and found that the correlation between the control participants’ true BOLD signal and the BOLD signal of their yoked participant from the treatment group was very low (see supplementary methods).

### Data Processing and Analyses

#### MRI Acquisition Parameters

All scanning was performed on a Siemens TIM Trio 3T scanner using a standard 32-channel head coil. For the fMRI tasks we used a whole-brain multi-shot 3D echo-planar imaging (EPI) sequence(Lutti *et al.*, 2013) with TR = 1.2 s, TE = 30ms, excitation flip angle = 15°, bandwidth = 2604 Hz/Px. There were 60 slices per slab, acquired with sagittal orientation and anterior to posterior phase encoding. Image in-plane resolution was 64×64 and voxel size = 3×3×3 mm^3^. To allow fast whole-brain coverage we used GRAPPA parallel imaging in phase encoding and partition encoding direction with 2×3 acceleration. Quantitative Multi-Parameter Maps(Draganski *et al.*, 2011; Weiskopf *et al.*, 2013; Callaghan *et al.*, 2014) and diffusion weighted imaging (DWI) scans were also acquired during the baseline and three follow-up sessions. The acquisition details are included in the supplementary methods. Because we did not find any significant differences in the fMRI data to suggest successful training and transfer, we did not proceed with the statistical analysis of the MPMs and DWI images.

#### Real-time fMRI Setup

For the NFT sessions the EPI volumes were exported using Ice and Gadgetron(Hansen and Sørensen, 2013). In-house scripts created using Gadgetron and MATLAB (Mathworks) were used to reconstruct the 3D EPI data using SENSE(Pruessmann *et al.*, 1999) such that they could be read in near real-time by TBV to produce the target ROI time-series. There was a small delay at the start of each run to enable MATLAB to start, but after about 15s both the MRI scanner and the Gadgetron pipeline were fully in-synch with approximately 1s latency. To enable both systems to synchronize we introduced a delay of 18 volumes at the start of each run. During that time participants viewed a white cross on a black background followed by a count-down (from 10 to 1) until the NFT paradigm started. In-house MATLAB scripts were used to process the ROI time-series and record participants’ responses, breathing and heart rate. For the NFT runs, the ROI signal was regressed against head motion traces and physiological noise from respiration(Birn *et al.*, 2008) and cardiac rhythm using RETROICOR(Glover *et al.*, 2000). The “cleaned” signal was then processed by in-house MATLAB scripts using Cogent toolbox (http://www.vislab.ucl.ac.uk/cogent_2000.php) to calculate and present the feedback to the participant. The computer setup in the scanner is shown in Supplementary Figure 1B.

#### Data Processing

All statistical analyses were performed after extensive quality control and offline pre-processing of the fMRI data. Supplementary Figures 3 and 4 show the evoked response patterns and average correlation coefficients respectively from the real-time processing pipeline. No statistical analyses were performed on these data, but are presented here for completeness. Statistical Parametric Mapping SPM12 (Wellcome Trust Centre for Neuroimaging, London) was used for offline pre-processing of the fMRI data. The first 3 volumes were removed from all fMRI time series apart from the NFT runs, where we removed the first 18 volumes. The images were then corrected for head-motion with rigid-body realignment using a 2-step approach.

For the ROI analyses the re-aligned images were smoothed in native space using an isotropic 8mm FWHM Gaussian smoothing kernel. First-level, within-subject models included the condition of interest and noise regressors. We used 2 regressors modelling the upregulation and response (feedback blocks in the case of connectivity NFT) blocks for the baseline, NFT and transfer runs, and 1 regressor modelling the fist clenching blocks for the localiser runs. The baseline condition was modelled implicitly. In addition, first-level models included 6 head motion parameter regressors produced by SPM and extracted from the PMCS (where applicable) with their temporal derivatives, the quadratic expansions of the movement parameters and their derivatives(Friston *et al.*, 1996; Ciric *et al.*, 2017), spike regressors (see supplementary materials(Lemieux *et al.*, 2007)), as well as 13 physiological noise regressors modelling the heart rate using RETROICOR and respiration(Glover *et al.*, 2000; Birn *et al.*, 2008; Hutton *et al.*, 2011; Misaki *et al.*, 2015). Temporal autocorrelation was modelled using SPM’s first-level autoregressive process (AR(1)) and a high-pass filter with 128s cutoff.

For the activity NFT group, contrast values for upregulation vs baseline were extracted for the target ROI for each session and the highest 10% of t-values(Todd *et al.*, 2017) were used to calculate the average ROI value. For the connectivity NFT group, the time-series for the target ROIs (SMA and striatum) was extracted using a 6mm sphere centred on the peak for upregulation vs baseline across all runs. The Pearson’s correlation coefficient of the time-series between the two ROIs within the upregulation periods was then calculated and transformed into Fisher z-scores.

#### Statistical Analyses

Because the two NFT approaches use a different feedback measure, i.e. contrast estimates in the case of activity NFT and correlation coefficients in the case of connectivity NFT, we converted the activity and connectivity estimates to standardized scores in order to be able to compare them directly. In more detail, the SMA activity estimates and Fisher transformed SMA-striatum correlation coefficients were converted into z-scores using the mean and standard deviation from the baseline fMRI runs in the activity and connectivity NFT groups respectively. The standardized activity and connectivity NFT target estimates were then used as outcomes in repeated-measures ANCOVAs with group (treatment vs control), NFT type (activity vs connectivity), session and their interactions as fixed effects. Baseline level of the NFT target and its interaction with NFT type were included as covariates in all analyses to increase model sensitivity(Dimitrov and Rumrill, 2003). Session was modelled as a repeated factor within subjects. The primary endpoints for this study were NFT learning and near transfer. For the analyses testing for learning, session was modelled as a numerical factor, increasing from 0 to 3, to test for a linear increase across the training sessions. For the analyses testing for transfer effects, session was modelled as a categorical factor. Intersession covariance was modelled using heterogeneous compound symmetry (CSH), as this gave a reasonable approximation of the observed within-subject covariance while using minimal degrees of freedom. Model residuals were visually inspected using Q-Q plots and histograms for outliers and to ensure residuals meet normality assumptions. We used SAS 9.4 mixed approach to estimate the ANCOVAs. Because we used standardized measures for all the analyses, the model estimates provided are in units of standard deviation.

For the exploratory ROI analyses testing the relationship between learning and self-regulation ability in the NFT target levels with behavioural change we used between-group ANCOVAs. To extract the learning slope per participant we re-fitted the repeated-measure ANCOVA described above specifying random slope and intercept. As a measure of self-regulation ability we used the difference in NFT target level at the first and third follow-up sessions during upregulation without feedback compared to baseline. Other factors included in these models were group, NFT type and their interactions, as well as the baseline measure of cognitive and psychomotor function using the composite score. The dependent variable was the composite score at the first and third follow-up session. All tests were two-tailed and the alpha-level used to determine significance was p < 0.05.

### Data Availability

All data are available from the authors. Raw data cannot become publicly available due to lack of consent from the study participants.

## Results

### Learning Effects: Increase across training sessions

To examine differences in NFT learning between the treatment and control groups and the two different types of NFT we used repeated-measures ANCOVA testing for between group differences across all NFT sessions, as well as a linear increase in the target NFT measure(Hellrung *et al.*, 2018) across sessions. The dependent variable was the standardized NFT target estimates and the model included as factors session (modelled as a continuous variable), group (treatment vs control), NFT type (activity vs connectivity) and all their interactions. The model was also adjusted for baseline NFT target levels and its interaction with NFT type. The main effect of group tests for differences in the change from baseline between the treatment and control groups across all visits, whereas the group by session tests for differences in learning slope between groups. To test the main effect of group across all training sessions we used least square mean testing and compared the NFT target estimates across all NFT sessions between treatment and control groups.

There was no evidence for a difference between the activity and connectivity treatment groups in learning slope (p > 0.6 for both group by NFT type and group by NFT by session interactions. Linear increase across sessions estimate 95% CI: activity treatment = 0.078 (−0.190, 0.345); connectivity treatment = −0.056 (−0.323, 0.211); activity control = −0.057 (−0.324, 0.210); connectivity control = - 0.519 (−0.319, 0.215); see Figure 2C). However, there was a significant main effect of group, where the treatment group had greater NFT target levels overall compared to the control group across all visits (t(29.1) = 2.79, p = 0.009. Estimate 95% CI of group difference across all sessions: 0.816 (0.22, 1.41). Because NFT target levels were standardized the unit of the estimates is standard deviations; see Figure 2D). At baseline, there was no difference in NFT target levels between the groups (F(1, 28) = 0.20, p = 0.655; see supplementary materials), our findings therefore suggest that the effects of receiving neurofeedback occurred within the first training session and were stable across training sessions.

### Near Transfer: Upregulation without feedback

After the four training sessions, participants returned for three follow-up sessions. During those sessions we examined the ability of the participants to self-regulate the NFT target levels without receiving any feedback (i.e. near transfer). Similar to our previous analyses examining learning effects, we used a repeated-measures ANCOVA with factors group, session (modelled as a categorical factor in this model), NFT type and their interactions, adjusting for the baseline NFT target levels and its interaction with NFT type. The dependent variable was the standardized NFT target contrast estimates (upregulation without feedback compared to no upregulation). In this case, we did not hypothesize any difference between sessions and expected that transfer effects would remain stable for the three follow-up visits. Therefore, the effects of interest were the main effect of group (treatment vs control), which tested between-group differences in the increase of the NFT target levels across all follow-up visits from baseline, and the group by NFT type interaction (treatment vs control by activity vs connectivity NFT).

Although the treatment group increased NFT target levels at the follow-up visits compared to baseline by 0.9 standard deviations (estimate 95% CI increase from the baseline session: treatment group = 0.929 (0.341, 1.518); control group = 0.186 (−0.407, 0.778)), this increase was not significantly different from the control group (F(1, 26.4) = 3.27, p = 0.082. Estimate 95% CI of the group difference across all sessions = 0.74 (−0.10, 1.59); see Figure 3A). The connectivity treatment group was the only group able to increase its NFT target levels at follow-up compared to baseline (estimate 95% CI increase from baseline across all follow-up sessions: connectivity treatment = 1.226 (0.389, 2.063); activity treatment = 0.633 (−0.195, 1.460); activity control = 0.322 (−0.505, 1.149); connectivity control = 0.0492 (−0.799, 0.897); Figure 3A). However, the interaction between NFT type and group was not significant (F(1, 26.4) = 1.11, p = 0.30). There were no other significant effects or interactions (all p > 0.29). Our results suggest that although there is some evidence regarding a near transfer effect in the treatment group, particularly in the connectivity treatment group, it is weak and not significantly better than the control group.

**Figure 3:**
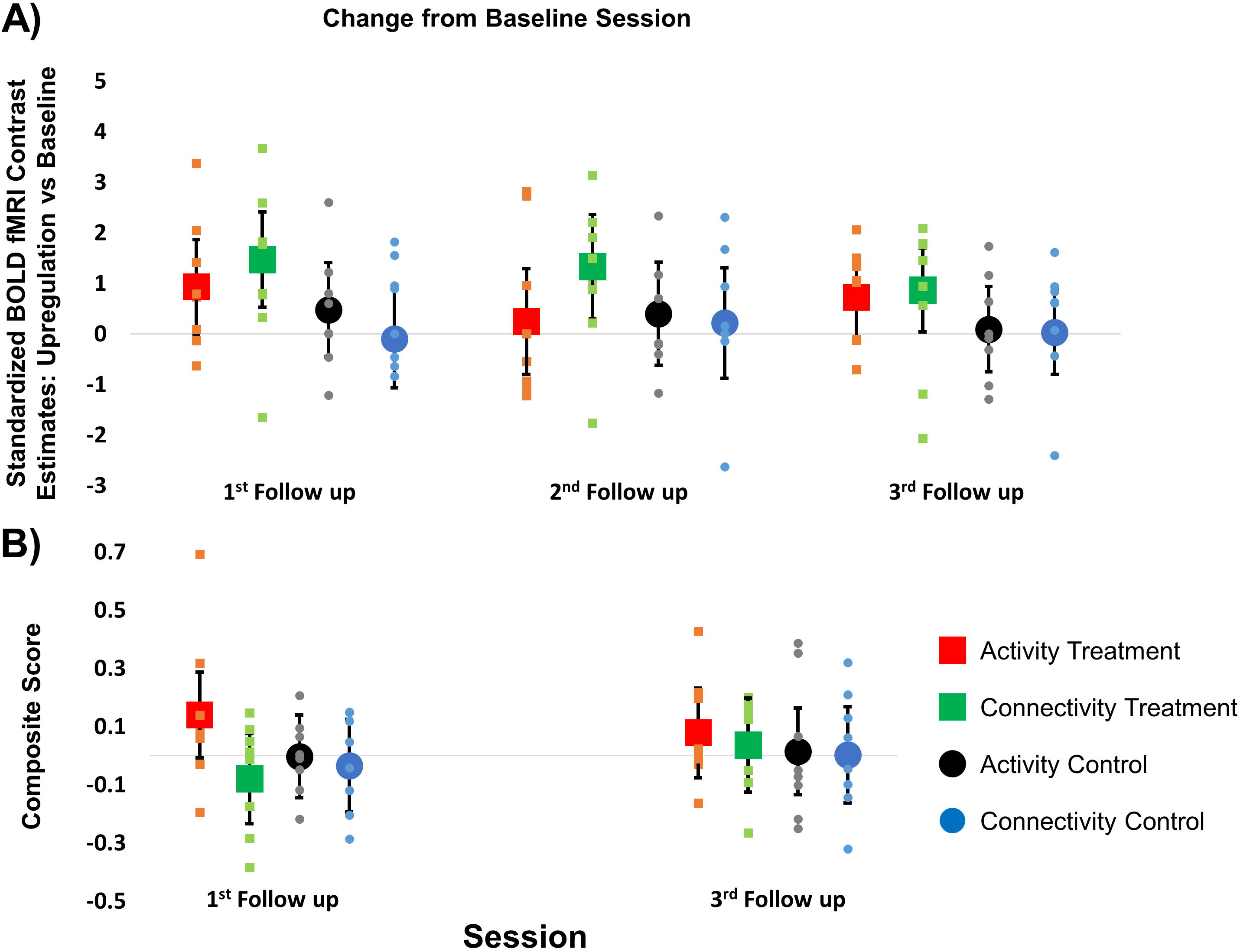
Near and far transfer effects. (A) Dot plots show the change in NFT target levels from baseline across the 3 follow-up sessions for the four subgroups: activity treatment group (red circles), connectivity treatment group (green circles), activity control group (black squares) and connectivity control group (blue squares). (B) Dot plots show the change in the behavioural composite score from baseline across the 2 follow-up sessions for the four subgroups (same colour coding as above). The horizontal gray lines in both plots show the baseline, data points above this line represent an increase compared to baseline. The small squares and circles show the individual data points, whereas the larger squares and circles show the adjusted mean group effects. Error bars are 95% CI.

### Far Transfer: Cognitive and psychomotor performance

To assess the effect that NFT had on participants’ performance in tasks unrelated to the training (far transfer), we examined change from baseline after training in the composite score comprising of measures previously shown to be sensitive to HD progression (*Tabrizi* et al., *2011*; *Papoutsi* et al., *2018*). We performed a similar mixed linear model analysis to the one described in the near transfer section above. The cognitive composite score was the dependent variable and the model was adjusted for the baseline level of the cognitive composite score. The effects of interest were the main effect of group and the group by NFT type interaction, which test for between group differences in change from baseline across all the two follow-up sessions.

Although the difference between treatment and control groups is in favour of the treatment group (estimate 95% CI increase from baseline: treatment = 0.044 (−0.059, 0.146); control = −0.005 (−0.102, 0.091)), it is not significant (F(1, 27) = 0.63, p = 0.435) and the magnitude of the change is small (estimate 95% CI difference = 0.049 (−0.077, 0.175) standard deviations). There was also no evidence for a difference between the activity and connectivity treatment groups (group by NFT type interaction F(1, 27) = 0.76, p = 0.39. Estimate 95% CI increase from baseline: activity treatment group = 0.108 (−0.023, 0.240); connectivity treatment = −0.022 (−0.162, 0.119); activity control = 0.006 (−0.120, 0.131); connectivity control = −0.016 (−0.161, 0.130); see Figure 3B). There were no other significant effects or interactions (all p > 0.18). Our results therefore do not provide any evidence for a significant far transfer effect of NFT.

A detailed description of the average change in the individual scores that comprised the composite score is presented in the supplementary materials and can provide more insight on the magnitude of change in the individual tests (see supplementary Figure 5 and supplementary Table 3).

### Regression with Behavioural Change

As an exploratory analysis we examined whether NFT-related measures, specifically learning slope and/or self-regulation ability (near transfer), predict improvement in the cognitive composite score. If successful upregulation has an effect on behaviour than we should see a relationship between increasing ones NFT target levels and improvement in behaviour after training.

We first tested the relationship between training slope, i.e. change from the first to the last NFT training session, and behavioural performance at the first follow-up session, i.e. within two weeks from the end of training. The NFT learning slope for each participant was extracted from a random slope and random intercept mixed linear model testing for linear increase in NFT target levels across visits. To test the effect of NFT learning on behaviour, we used an ANCOVA with factors NFT target level slope (linear increase across NFT sessions), group (treatment vs control), NF type (activity vs connectivity) and their interactions. The model was also adjusted for the cognitive score at baseline and the dependent variable was the composite score at the first follow-up. The effects of interest were the main effect of learning slope and its interactions with group and NFT type. The main effect of learning slope tests whether there is a relationship between increase in NFT slope across training sessions and improvement in behaviour at follow-up across all groups. The interactions between learning slope and group (and NFT type), test whether there is a difference in the relationship between improvement in behaviour across the different groups.

The relationship between NFT learning slope across all groups and change in the composite score was not significant (F(1, 23) = 2.67, p = 0.12) and negative (estimate 95% CI of change in the composite score per unit increase in slope = −0.59 (−1.35, 0.16)). There was also no significant difference between treatment and control groups in the relationship between NFT slope and composite score change (F(1, 23) = 0.89, p = 0.36; estimate 95% CI of the between group difference = 0.71 (−0.74, 2.15)), or evidence of a positive relationship in the treatment group (estimate 95% CI of change in the composite score per unit increase in slope for the treatment group = −0.24 (−1.24, 0.78); control group = −0.94 (−2.00, 0.12); see Figure 4A). All other effects and interactions were also non-significant (all p > 0.1).

**Figure 4:**
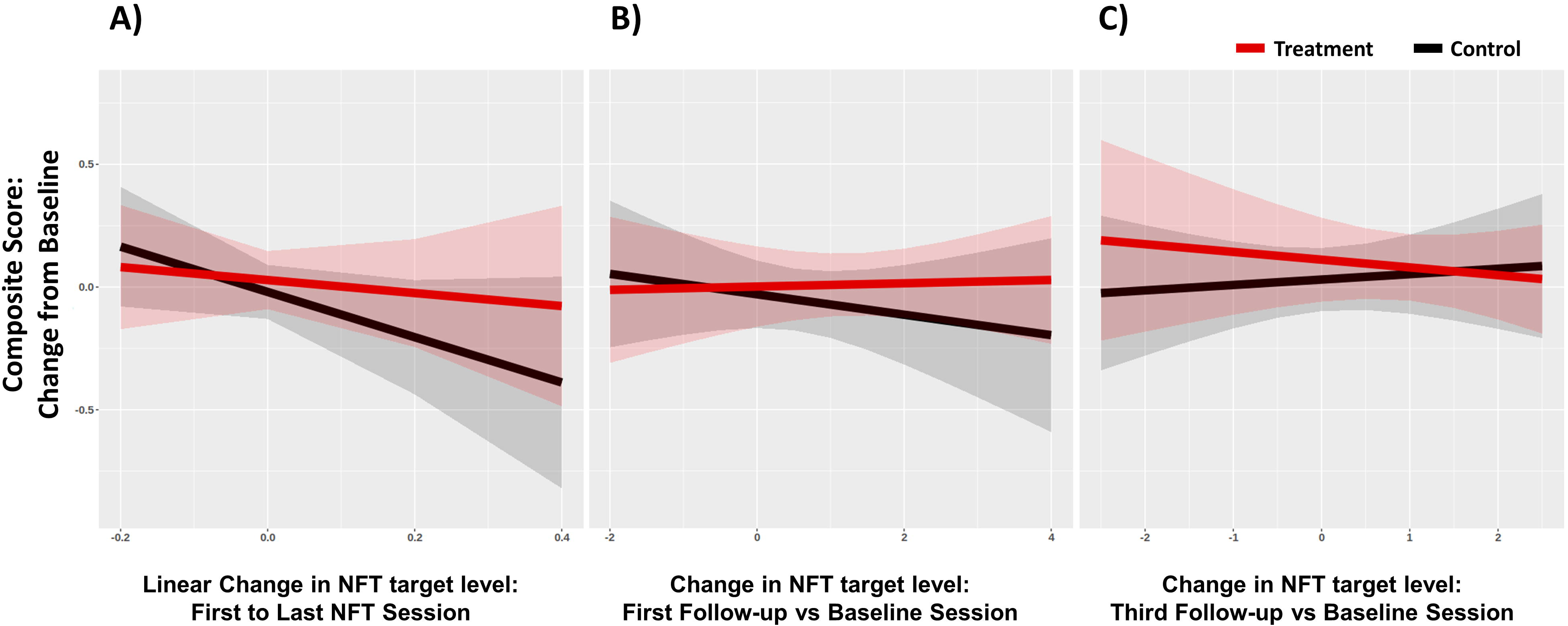
Relationship between change in the composite score and change in NFT target levels. (A) Regression lines plot the relationship between change in the composite score at the first follow-up from baseline and change in NFT target levels from the first to the last NFT training visit adjusted for baseline levels. Shown in (B) is the relationship between change in the composite score at the first follow-up from baseline and change in NFT target levels at the first follow-up session compared to baseline. Shown in (C) is the relationship between the same measures as in (B), but for the third follow-up. Regression lines and 95%CI for the treatment (red) and sham control (black) groups are averaged across both NFT type groups.

We then tested the relationship between volitional NFT-target upregulation ability and improvement in the composite score at follow-up (first and third follow-up separately). We used ANCOVA with factors NFT target level estimate (the difference between NFT target contrast estimates at each follow-up vs baseline), group (treatment vs control), NF type (activity vs connectivity), and their interactions. The model was adjusted for baseline performance in the composite score and the dependent variable was the composite score at the first and third follow-up. The effects of interest were the relationship between behavioural change at follow-up and self-regulation ability, as well as the interaction with group and NFT type.

The relationship between NFT-target upregulation ability across all groups and change in the composite score was not significant for either of the follow-ups (first follow-up: F(1, 22) = 0.28, p = 0.60; third follow-up: F(1, 22) = 0.02, p = 0.90) and almost zero in both cases (estimate 95% CI of change in the composite score per unit increase in slope for the first follow-up = −0.02 (−0.09, 0.05) and for the third follow-up = −0.01 (−0.08, 0.07)). There was also no significant difference between treatment and control groups in the relationship between NFT slope and composite score change (first follow-up: F(1, 22) = 0.56, p = 0.46; estimate 95% CI of the between group difference = 0.08 (−0.09, 0.25); third follow-up: F(1, 22) = 0.48, p = 0.50; estimate 95% CI of the between group difference = 0.03 (−0.16, 0.22); see Figure 4B and 4C). All other effects and interactions were also non-significant (all p > 0.1).

## Discussion

The present proof-of-principle study examined the use of NFT for the treatment of cognitive and psychomotor impairment in HD patients. For this purpose, we used two different NFT approaches (activity and connectivity) in a single-blind, RCT study, with yoked sham NFT control groups, an intensive training protocol consisting of 16 NFT trials over 4 sessions, optimized real-time fMRI acquisition protocol and using objective, a-priori defined, measures of cognitive and psychomotor function. This enabled us to collect rigorous evidence regarding the usefulness of NFT in treating cognitive and psychomotor symptoms in HD. We found strong evidence of a difference between treatment and control groups during the NFT sessions, such that participants in the treatment group increased the levels of the NFT target more than participants in the control group, when receiving NFT. However, evidence regarding the ability of the participants to volitionally upregulate their NFT target levels after training was weak. It is therefore unclear whether participants learned to regulate their brain activity and were able to apply the learning in the absence of NFT. We also did not find any evidence of improvement in cognitive and psychomotor function after training in the treatment group, or of a relationship between NFT-learning and change in cognitive and psychomotor function.

In more detail, we found a significant difference between treatment and control groups in terms of the increase of their NFT target levels from baseline. Participants in the treatment group increased their activity and connectivity levels from baseline by 0.74 standard deviations more than participants in the sham control group. This difference was present from the first training session until the last and we did not observe any further increase in the subsequent training sessions. This finding is in agreement with previous studies which have shown that participants can learn to regulate the target NFT levels within one visit(Hellrung *et al.*, 2018; Kohl *et al.*, 2019).

It is important to note that our study was single-blind, it is therefore possible that the difference observed between treatment and control groups could have been because of unconscious researcher bias. We believe that this is unlikely, since the participants were in the MRI scanner during NFT and they had minimal contact with the researchers. In addition, if they were such effects, we would expect that they would have been more pronounced during cognitive and psychomotor testing, during which the researchers had longer contact with the participants. However, we did not find any evidence for a difference between the two groups, treatment and controls. Therefore, we believe that the measured difference between the two groups during NFT training reflects the effect of providing feedback to the participants on their NFT levels and them adjusting their behaviour accordingly.

After NFT we tested participants’ ability to increase the levels of the NFT target without receiving neurofeedback. This way we can to test whether participants have truly learned to regulate the levels of the NFT target and are therefore able to upregulate without receiving NFT. Although the treatment group increased their NFT target levels from baseline at follow-up by 0.9 standard deviations, this increase was not significantly different from the control group. Our results therefore suggest that although there is some evidence regarding a near transfer effect in the treatment group, particularly in the connectivity treatment group, it is weak and at present ambiguous.

Furthermore, we did not find any evidence of improvement in cognitive and psychomotor function after NFT. To measure cognitive and psychomotor function we used a composite score comprised of a-priori identified, objective measures of cognitive and psychomotor function, sensitive to HD progression. Although the between-group difference in the change from baseline was in favour of the treatment group, the magnitude of the improvement was very small and clinically non-significant representing a change of 0.05 standard deviations in the composite score. We also did not find any evidence that change in NFT-related measures, specifically NFT learning slope and ability to self-regulate, related to change in behaviour. It is therefore doubtful whether NFT, even if it was successful, could lead to improved cognitive and psychomotor function in HD.

Taken together our findings are in agreement with recent studies showing a lack of clinical benefit of NFT using objective measures of brain activity and behaviour, despite evidence for differences between treatment and control groups during NFT(Schabus *et al.*, 2017). Therefore, even if it is possible to train people to self-regulate using NFT, at present it is not clear whether this method, on its own, will lead to a clinical benefit.

The failure to find reliable evidence of clinical benefit could be because we did not target the right regions or connections. In this study we used two different NFT targets, SMA activity and SMA-striatum connectivity. The latter was selected based on our previous work which showed that improvement in cognitive and psychomotor function predicted increased SMA-striatum connectivity across NFT sessions(Papoutsi *et al.*, 2018). In the present study, despite using the same cognitive and psychomotor measures as in our previous study and targeting the networks identified in our previous study, we did not find any evidence to suggest that SMA-striatum connectivity NFT relates to improvements in cognitive and psychomotor function. Therefore, we were not able to replicate this finding from our previous work. It is possible that other regions and connections, such as the left inferior parietal lobe, which have been implicated in neural compensation in HD(Klöppel *et al.*, 2015), could have been more appropriate. This remains to be tested.

Finally, in our study we did not find strong evidence to suggest a difference between the two NFT methods, activity and connectivity. The connectivity treatment group was able to increase the NFT target levels at follow-up compared to baseline, however, there were no significant group differences. Although targeting SMA-striatal connectivity is theoretically motivated by knowledge regarding the disease mechanism and was identified as a potential target in our previous study(Papoutsi *et al.*, 2018), we did not find any reliable evidence that it was better or worse than activity NFT. Both activity(Young *et al.*, 2017; Hellrung *et al.*, 2018) and connectivity(Megumi *et al.*, 2015; Ramot *et al.*, 2017; Yamashita *et al.*, 2017) NFT have been used successfully in other studies, suggesting that both methods are effective.

A limitation of our study is that we could not dissociate the use of connectivity NFT from the use of intermittent feedback. In our study, feedback was provided continuously in near real-time in the activity NFT group, whereas in the case of connectivity NFT, correlations were computed over 30secs and the feedback was presented intermittently at the end of the upregulation block. Therefore, the two elements, frequency of feedback presentation and NFT type, were intertwined and could not be separated. A previous study comparing continuous vs intermittent feedback using percent signal change in the amygdala in healthy young adults showed that participants were able to learn to increase the target NFT levels using both approaches, although intermittent feedback was more effective than continuous in that study(Hellrung *et al.*, 2018). In our study we did not find any evidence for a difference between the two approaches.

To conclude, in the present study we compared two different NFT approaches in HD, SMA activity and SMA – left striatum connectivity NFT against sham NFT control groups, in terms of learning and transfer. We used an RCT design and an intense, optimized real-time fMRI NFT protocol, to ensure that we can acquire rigorous evidence regarding the role of real-time NFT in HD. Our findings support previous claims that using NFT participants can be guided to increase their levels of cortical activity and cortico-striatal connectivity using real-time fMRI NFT. However, evidence regarding the transfer of learning to volitional control of brain activity and behaviour are currently weak. The clinical usefulness of real-time fMRI NFT in HD is therefore doubtful and at present we cannot recommend the use of NFT real-time fMRI as an intervention to improve cognitive and psychomotor symptoms in HD.

## Acknowledgments

We are very grateful to the study participants and their families for their support. We would also like to thank the imaging support team at the Wellcome Centre for Human Neuroimaging and our funders.

## Authors’ Roles

M.P., D.L, N.W., G.R. and S.J.T designed the study and prepared the manuscript; M.P. performed neuropsychological testing, MRI data acquisition, data processing and statistical analysis; J.M. implemented the real-time fMRI pipeline and PMCS; O.J. programmed the gadgetron pipeline; T.I. and S.P. assisted with data collection and patient recruitment; N.H. and E.P. created the mouth pieces for the PMCS; D.L. supported the statistical analysis. R.R. supported the processing of the Q-Motor data. All authors discussed results and commented on the manuscript. G.R. and S.J.T. contributed equally to the study.

