## Supplementary for "Activity or Connectivity? Evaluating neurofeedback training in Huntington’s disease"

### Supplementary Materials

*Sample size estimation*

The present study is the first randomized controlled neurofeedback (NFT) study in HD, since our previous pilot NFT study in HD did not include a control group[1]. Therefore our sample size of 32 participants was based on a sham-controlled NFT study on healthy young adults that included pre- and post-raining comparisons in near-transfer[2]. This study from Yoo et al.[2] found large effect sizes with 11 participants per group. In more detail we estimated a Cohen's d effect size of 1.65 and 1.90 (immediate and two-week post-training follow-up respectively) for differences between the treatment and sham control group in near transfer effects (follow-up vs baseline). With 8 participants per group we would therefore have 80% power with 5% type I two-sided error to detect larger than 1.51 ES.

*Prospective Motion Correction*

Many of the patients in our cohort had large involuntary movements (chorea) that could interfere with the scanning and result in many scan fails. Therefore, to improve quality of the scans and minimize data loss because of head motion, we employed a prospective motion correction system (PMCS). The system we used consists of an optical camera mounted inside the MRI scanner bore which tracks the motion of a passive marker[3]. Information about the position of the marker is then sent to the scanner to dynamically update the imaging FoV. Details of the system we used and how it works can be found in previous papers[4,5]. The PMCS display in the scanner room and setup is shown in Supplementary Figure 1. Because the PMCS has a time lag of 23ms we improved the system by developing a GLM-based algorithm that performed head motion prediction. This allowed faster and smoother motion correction.

A small retainer was used to hold the marker for the PMCS. The retainer was custom-made for each participant by an orthodontic technician (EP) using the dental impressions acquired during the screening visit. The retainer was moulded to a plastic Lego© chair (www.lego.co.uk) and attached to other Lego© pieces such that the marker was positioned in the centre of the PMCS field of view, when the participant's head was moved inside the MRI scanner. An example of the setup is shown in Supplementary Figure 1A. This setup for the PMC marker is light and easy to use. It allows participants to speak and therefore report any problems or discomfort during the scanning session. Nylon screws were used to ensure that all moving parts near the participant's mouth were safely

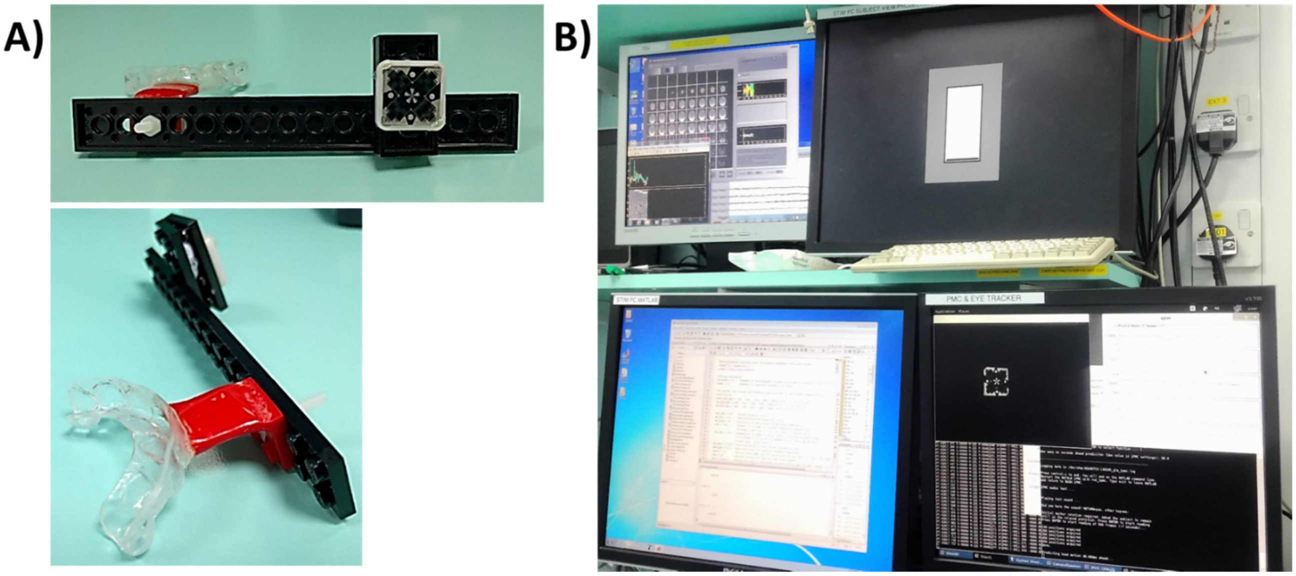

*Supplementary Figure 1:PMCS and experimental Setup. (A) A small retainer was attached to the PMCS marker with Lego© pieces. The retainer clipped to the front of the teeth. (B) The setup in the scanner control room that was used to monitor and deliver stimuli in the MRI scanner. The top left screen shows the TBV and custom Matlab setup used to record BOLD fMRI signal in near real-time. The top right screen displays the screen that the participant views in the scanner. The bottom left screen shows the Cogent script that was used to compute and present the neurofeedback. The bottom right screen shown the PMCS monitoring system. The location of the marker with respect to the camera's field of view is shown in the top left quarter of the screen.*

attached and would not disconnect during a sudden head movement and e.g. fall inside a participant's mouth. Use of the PMCS was not compulsory in order not to hinder recruitment to the study and 12 out of 32 participants did not use it. *Supplementary Table 1* shows the number of participants that used the PMCS across groups. The same sequences were used for all participants irrespective of whether they used the PMCS or not. When PMCS was used PMCS XPACE Motion Correction (MoCo) was enabled in the sequence.

*Participant randomization*

The UHDRS TMS was selected as the basis of the randomization because it is a marker of clinical disease impairment. Randomization was performed by the study's statistician, DL. We used a TMS cutoff of 10 to divide patients into high and low TMS groups (low group <= 10) and randomized participants so that each experimental group (treatment and control) had comparable number of participants from each TMS group. A constraint in randomization was imposed by the fact that participants in the sham control group were yoked to participants in the treatment group (1:1) and received feedback based on the activity from the treatment group. This meant that the first participant for the activity and connectivity groups had to be allocated to the treatment group and the last participant was allocated to the control group. The randomization to groups was done using a random number generator in R. This was a single-blind study, so participants did not know which group they belonged to for the duration of the study.

*Data quality check: spike regressors*

Six motion parameters were generated for each fMRI run from SPM12. In participants who used the PMCS the motion parameters represented residual and not actual head movement, i.e. they reflected motion that could not be fully corrected by the PMCS. For this reason, in order to identify and exclude bad scans we did not use the motion parameters, e.g. by examining scan-to-scan motion. Instead we used the DVARS[6] approach developed by Afyouni and Nichols[7]. Volumes were identified as bad if the change in DVARS was greater than 20% and were added as separate regressors to the first-level models. Blocks where more than half of the volumes were de-weighted were excluded.

Similar to our previous study we also monitored overt hand movements using pneumatic tubes[1] and inspected the upregulation and rest blocks (details are provided in the task compliance section below). If a response was detected, the block was excluded from the analysis. Runs with less than 2 blocks remaining in the analyses were excluded completely. If the data from all runs within a session were excluded, then this participant was excluded from those analyses that included this session only. Supplementary Table 3 shows the number of participants whose data were used in the analyses for each session.

*Task Compliance*

Prior to scanning, all participants were instructed to refrain from making any overt movements and only use mental strategies, such as motor imagery, in order to increase the levels of the NFT target (represented by the height of the red bar). Compliance was monitored using pneumatic tubes similar to our previous study[1]. Specifically, during the NFT and transfer runs the participants' hand movements were monitored during the localiser and neurofeedback training runs using pneumatic tubes connected to a pressure sensor which were taped along the participants' palms. This setup enabled the detection of gross limb movements, e.g. flexing one's fingers or squeezing one's fist, during the scanning session and ensured task compliance. If a participant made overt movements during any of the blocks, they were reminded that they should not do them and these blocks were excluded during post-processing from the analysis (see section on offline fMRI analyses below).

*Baseline Levels of the NFT Target Measure*

The baseline session included a motor imagery task used to calculate the levels of the NFT target measure prior to NFT (see Methods section in the main manuscript for details on the experimental design). For the activity NFT group we measured the BOLD fMRI signal from the SMA during motor imagery compared to the control condition, whereas for the connectivity NFT group it was the time-series correlation between the SMA and the left striatum during upregulation (SMA-striatum connectivity). Because participants were randomized we did not expect any systematic differences between the groups in terms of the level of their brain activity and connectivity prior to NFT. To confirm this we ran a two-way ANOVA with factors group (treatment vs control), NFT type (activity vs connectivity), as well as their interaction. There were no significant differences (Group: $F(1, 28) = 0.20$, $p = 0.655$; NFT type: $F(1, 28) = 0$, $p = 1.000$; NFT type by Group: $F(1, 28) = 1.11$, $p = 0.302$).

*Sham Control Group Confirmatory Analyses*

To confirm that control participants received feedback that was not contingent to their own brain activity, we examined the correlation between the brain activity of the control participants and their yoked treatment participants. For the activity NFT participants, we correlated the SMA ROI values recorded in real-time at every volume between the matched treatment and control participants. The Pearson's correlation coefficients were calculated per block and averaged across all blocks. These ranged between -0.003 to 0.099 for each pair of participants. We also counted the number of blocks that had a Pearson's $r > 0.381$, which for $n = 25$ timepoints corresponds to $p < 0.05$ uncorrected for multiple comparisons. We used this as an indication of the percent of blocks where participants in the control group would have received feedback contingent to their own brain activity. Out of 80 blocks in total the number of blocks for each pair of participants with r greater than 0.381 ranged between 2 and 17, i.e. 2.5% - 21% of the blocks. The correlation between the NFT signals of the yoked participants was therefore quite low, suggesting that the sham NFT approach worked.

In the case of the connectivity NFT, participants' feedback was presented only once at the end of each upregulation block and was based on the Pearson's correlation coefficients between the SMA and left Striatum time-series. To examine whether the sham feedback that was presented to the control group was related to their actual connectivity, we compared the correlation coefficients per block for each pair of treatment and control participants using the r.test function in R's psych package (R version 3.4.4; psych version 1.8.3.3). The number of blocks out of 80 that were significantly different ($p < 0.05$ uncorrected) ranged between 13 and 27, i.e. between 16.25% - 33.75% of the blocks. This percentage was higher than that for the activity NFT groups, but still relatively low.

*Structural MRI – acquisition parameters*

Quantitative Multi-Parameter Maps[8–10] were acquired during the baseline and three follow-up sessions. We acquired three spoiled multi-echo 3D fast low angle shot (FLASH) whole-brain volumes: (a) proton density (PD) weighted images with flip angle = 6°, TR = 25ms and with eight echoes TE=2.34ms, 4.64ms, 6.94ms, 9.24ms, 11.54ms, 13.84ms, 16.14ms and 18.44ms, (b) T1-weighted images with flip angle = 21°, TR = 25ms and with eight TE=2.34ms, 4.64ms, 6.94ms, 9.24ms, 11.54ms, 13.84ms, 16.14ms and 18.44ms, and (c) magnetisation transfer (MT) weighted images with flip-angle = 6°, TR = 25ms and with six TE=2.34ms, 4.64ms, 6.94ms, 9.24ms, 11.54ms and 13.84ms. To achieve magnetisation transfer weighting an off-resonance RF pulse was applied before non-selective excitation. All volumes had 0.8mm isotropic voxel resolution, a field of view (FoV) of 256x224 $mm^2$ and 224 slices. To shorten acquisition time, we used GRAPPA parallel imaging in phase encoding (anterior-posterior) and partition encoding (right-left) direction with 2x2 acceleration. Each scan lasted 7 mins. Prior to the acquisition of the MPMs, calibration data (B1 and B0 maps) were acquired to correct for inhomogeneities in the RF transmit field.

Three-shell DWI data were also acquired during the baseline and three follow-up sessions. The DWI protocol consisted of 3 b values of 2,000, 700, and 300s/mm2 with 64, 32, and 8 nonlinear diffusion-encoding directions, respectively; 14 b = 0s/mm2 images; voxel size = 2.0 × 2.0 × 2.0mm3; repetition time (TR) = 5,500 milliseconds; echo time (TE) = 98.0 milliseconds; 66 slices; multi-band factor = 2. To shorten acquisition time, we used GRAPPA parallel imaging in phase encoding (anterior-posterior) direction with acceleration factor = 2 and acquisition time = 11.11 minutes. To compensate for EPI geometric distortions an additional b = 0 s/mm2 image was acquired with opposite phase encoding directions. The geometric distortion was corrected with TOPUP and EDDY[11].

*References*

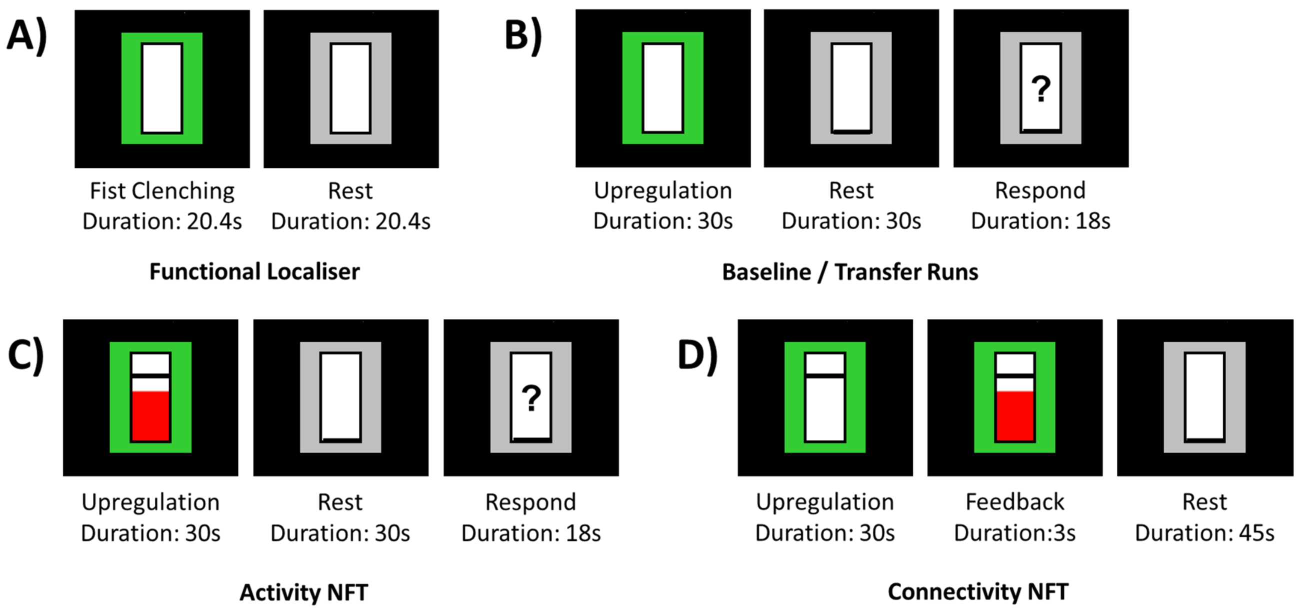


Supplementary Figure 2: Task design. Figure shows the display for the different blocks presented as part of (A) the functional localiser task, (B) the baseline and near transfer runs, (C) the activity-NFT training, and (D) the connectivity-NFT runs. A black line was set at 3/5 of the total bar height and acted as an additional reminder to the participants that they needed to increase the height of the red bar.

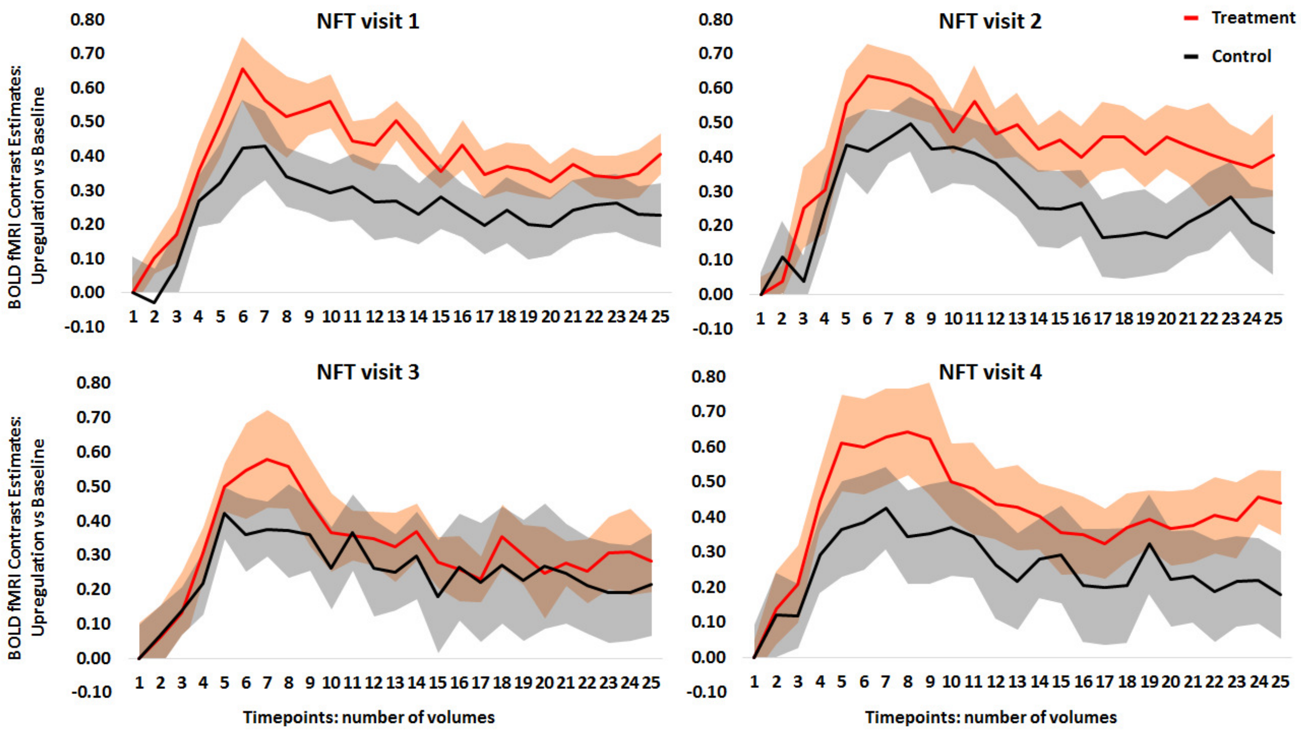


Supplementary Figure 3: Evoked response patterns. The group mean (SE) task evoked response pattern (ERPs) across all blocks per visit for the two activity NFT groups (treatment in red, control group in black). These data were measured and presented to the participants during the NFT visits and are presented here for completion. No statistical analyses were performed on these data. In all the analyses presented in the paper we used the data after extensive quality check and pre-processing as described in the methods sections.

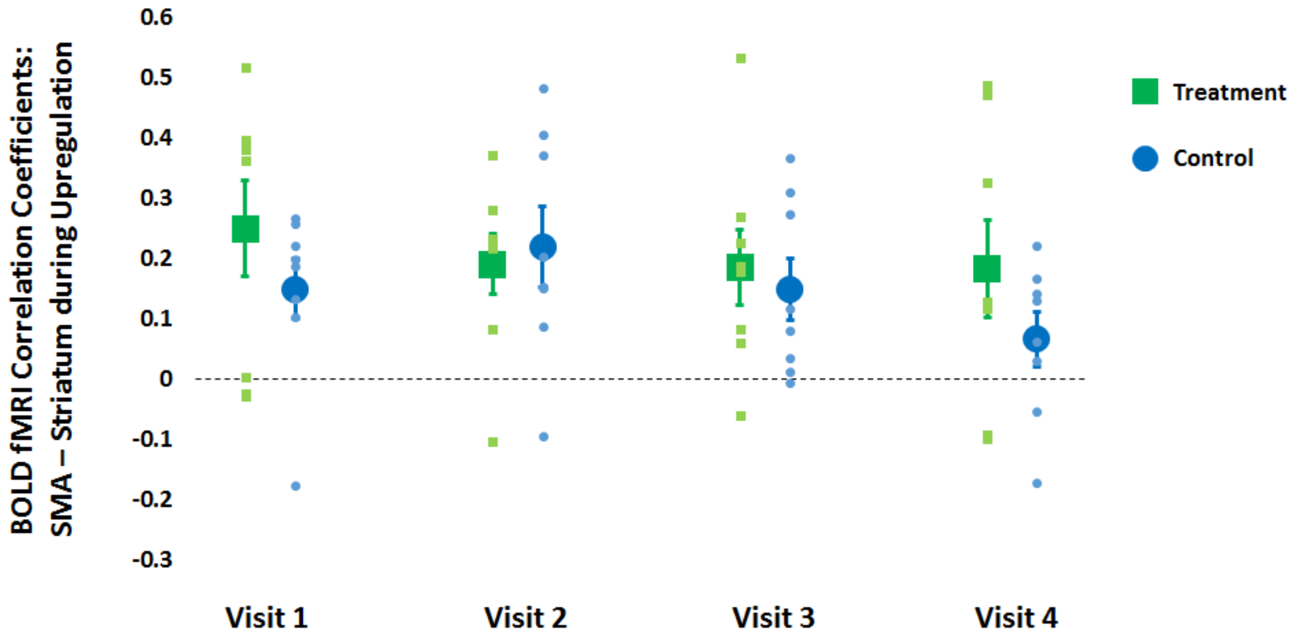


Supplementary Figure 4: Correlation Coefficients from the connectivity NFT groups (green for the treatment group, blue for the control group). Dot plots show for each NFT visit the mean (SE) correlation coefficients that were estimated in real-time for each block. These were presented as feedback to the participants, however no statistical analyses were performed on these data. In all the analyses presented in the paper we used the data after extensive quality check and pre-processing as described in the methods sections. The data are presented here for completion.

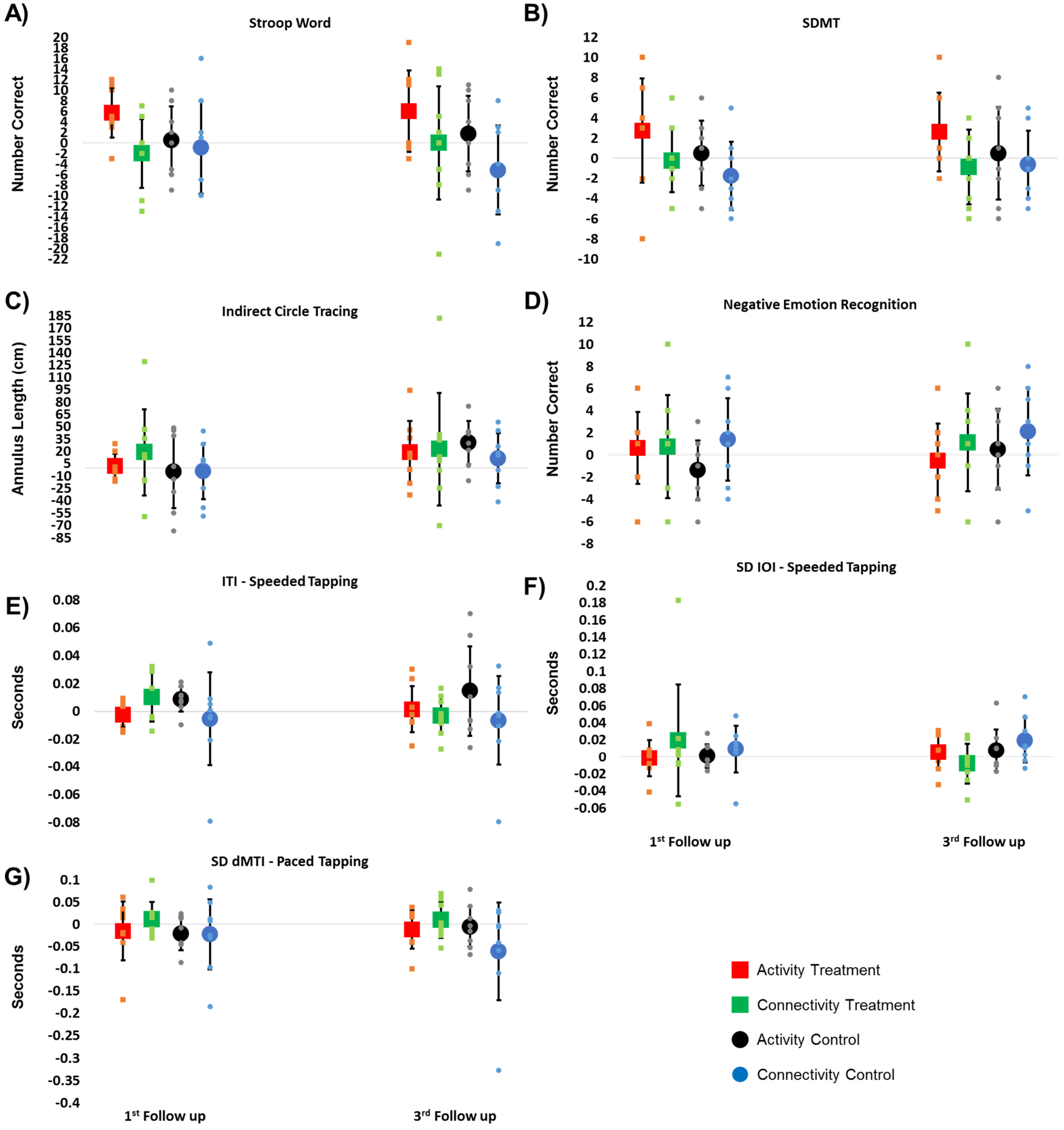


Supplementary Figure 5: Change from baseline in cognitive and psychomotor function. Dot plots show the change from baseline in the two follow-up sessions for each of the seven measures that were included in the behavioural composite score across the four subgroups: activity treatment group (red circles), connectivity treatment group (green circles), activity control group (black squares) and connectivity control group (blue squares). The values plotted are raw change scores. A value greater than zero (baseline) means that they improved in figures A-D. A value lower than zero means that they improved in figures E-G. SDMT = Symbol digit modalities test; ITI = inter-tap interval; SD IOI = standard deviation of inter-onset interval; standard deviation of mid-tap interval deviation from target rhythm.

*Supplementary Table 1: Information about the number of participants included in the imaging analyses.*

| | Number of participants using PMCS (out of 8) | Number of Participants included in Analyses | | |
|---|---|---|---|---|
| | | Baseline Session | NFT Sessions: 1 \| 2 \| 3 \| 4 | Transfer sessions: 1 \| 2 \| 3 |
| actNFT - Treatment | 6 | 8 | 8 \| 8 \| 8 \| 8 | 8 \| 7 \| 8 |
| actNFT - Control | 6 | 8 | 8 \| 8 \| 8 \| 8 | 8 \| 8 \| 7 |
| conNFT - Treatment | 3 | 8 | 8 \| 8 \| 8 \| 8 | 8 \| 8 \| 8 |
| conNFT - Control | 5 | 8 | 8 \| 8 \| 8 \| 8 | 7 \| 6 \| 8 |

*Supplementary Table 2: Information about session timing and completion. Three participants were not able to complete the second follow-up within the specified window because of MRI scanner unavailability or personal circumstances unrelated to the study. One participant could not complete the MRI session of the last follow-up because of the unavailability of the MRI scanner. For 7 participants the cognitive and psychomotor testing took place on a different day to the MRI scanning baseline or follow-up sessions to accommodate the participant's schedule or because of unavailability of the MRI scanner.*

| **Session** | **Number of Days Between Successive Sessions: Mean (SD)** | **Number of participants who completed the Session** |
|---|---|---|
| Screening | 0 | 32/32 |
| Baseline | 30.6 (36.9) | 32/32 |
| NFT 1 | 14.3 (11.6) | 32/32 |
| NFT 2 | 10.2 (4.9) | 32/32 |
| NFT 3 | 9.3 (5.5) | 32/32 |
| NFT 4 | 7.5 (2.7) | 32/32 |
| 1st post-training | 8.7 (3.4) | 32/32 |
| 2nd post-training | 25.7 (7.1) | 29/32 |
| 3rd post-training | 30.8 (7.4) | 32/32 (Behavioural), 31/32 (MRI) |

*Supplementary Table 3: Mean (SD) change in the cognitive and psychomotor tasks (raw scores) and the composite score from baseline at the first (gray rows) and third follow-up session (white rows). The "all" group category is all participants in the treatment and control groups collapsed across both NFT type groups. SDMT: Symbol digit modalities task; ND: non-dominant hand; ITI: inter-tap interval; IOI: inter-onset interval; dMTI: mid-tap interval deviation from target rhythm.*

| | **Treatment** | | | **Control** | | |
|---|---|---|---|---|---|---|
| | *Activity* | *Connectivity* | *All* | *Activity* | *Connectivity* | *All* |
| **Stroop Word:** *#Correct* | 5.75 (4.7) | -2 (6.5) | 1.88 (6.9) | 0.5 (6.4) | -0.88 (8.8) | -0.19 (7.7) |
| | 6 (7.7) | 0 (10.7) | 3 (9.8) | 1.75 (7.2) | -5.1 (8.5) | -1.7 (8.6) |
| **SDMT:** *#Correct* | 2.75 (5.2) | -0.25 (3.2) | 1.25 (4.5) | 0.5 (3.2) | -1.75 (3.4) | -0.63 (3.5) |
| | 2.63 (3.9) | -0.88 (3.7) | 0.88 (4.2) | 0.5 (4.6) | -0.63 (3.4) | -0.06 (4.1) |
| **Emotion Recognition:** *Negative, #Correct* | 0.63 (3.2) | 0.75 (4.6) | 0.69 (4.0) | -1.38 (2.7) | 1.38 (3.7) | 0 (3.5) |
| | -0.50 (3.3) | 1.13 (4.4) | 0.31 (4.0) | 0.50 (3.6) | 2.13 (4.0) | 1.31 (3.9) |
| **Indirect Circle Tracing:** *Annulus Length (cm)* | 2.1 (15.0) | 18.8 (52.1) | 10.4 (39.2) | -5.1 (44.3) | -4.1 (34.2) | -4.6 (39.6) |
| | 18.97 (37.6) | 22.6 (68.5) | 20.8 (55.3) | 30.5 (26.1) | 11.5 (30.7) | 21.0 (30.0) |
| **Speeded Tapping (ND):** *mean ITI (secs)* | -0.003 (0.009) | 0.010 (0.018) | 0.004 (0.015) | 0.009 (0.009) | -0.006 (0.034) | 0.002 (0.026) |
| | 0.001 (0.017) | -0.003 (0.014) | -0.001 (0.015) | 0.015 (0.032) | -0.007 (0.032) | 0.004 (0.034) |
| **Speeded Tapping (ND):** *SD IOI (secs)* | -0.001 (0.021) | 0.019 (0.066) | 0.009 (0.015) | 0.001 (0.014) | 0.001 (0.027) | 0.005 (0.022) |
| | 0.006 (0.021) | -0.008 (0.023) | -0.001 (0.023) | 0.008 (0.024) | 0.020 (0.026) | 0.014 (0.026) |
| **Paced Tapping:** *SD dMTI (secs)* | -0.015 (0.066) | 0.011 (0.039) | -0.002 (0.056) | -0.021 (0.037) | -0.023 (0.080) | -0.022 (0.062) |
| | -0.012 (0.043) | 0.010 (0.040) | -0.001 (0.043) | -0.005 (0.046) | -0.061 (0.110) | -0.033 (0.089) |
| **Composite Score** | 0.145 (0.25) | -0.070 (0.18) | 0.038 (0.24) | -0.003 (0.12) | -0.023 (0.16) | -0.013 (0.14) |
| | 0.084 (0.17) | 0.046 (0.15) | 0.065 (0.17) | 0.014 (0.23) | 0.014 (0.19) | 0.014 (0.21) |
